# Genome sequences of both organelles of the grapevine rootstock cultivar ‘Börner’

**DOI:** 10.1101/2020.03.18.996405

**Authors:** Bianca Frommer, Daniela Holtgräwe, Ludger Hausmann, Prisca Viehöver, Bruno Huettel, Reinhard Töpfer, Bernd Weisshaar

## Abstract

Genomic long reads of the interspecific grapevine rootstock cultivar ‘Börner’ (*Vitis riparia* GM183 x *Vitis cinerea* Arnold) were used to assemble its chloroplast and mitochondrion genome sequences. We annotated 133 chloroplast and 172 mitochondrial genes including the RNA-editing sites. The organellar genomes were maternally inherited to ‘Börner’ from *Vitis riparia*.

---

Long reads generated by the Single Molecule, Real Time (SMRT) DNA sequencing technology (Pacific Biosciences, PacBio) are one starting point for high-quality chloroplast (1, 2) and mitochondrion genome sequence assemblies. The cultivated grapevine *Vitis vinifera* is highly susceptible to pathogens. Resistant cultivars like the interspecific hybrid ‘Börner’ (*V.riparia* GM183 (mother plant) x *V.cinerea* Arnold (pollen donor)) are used as rootstocks for growing elite grapevine varieties. We assembled and annotated the chloroplast (cp_Boe) and mitochondrion (mt_Boe) genome sequences of ‘Börner’ from SMRT reads. All bioinformatic tools were applied with default parameters unless otherwise noted.

Genomic DNA was extracted from young leaves of ‘Börner’ (3) and sequenced on a Sequel I sequencer (1Mv3 SMRT cells, binding kit 3.0, sequencing chemistry v3.0, all from PacBio). Potential plastid or mitochondrial reads were filtered by blastn (BLAST 2.7.1+) searches (4) against plastid or mitochondrial sequences (RefSeq release 91). Criteria: read length above 500 nt, identity above 70%, query coverage above 30%. The 292,574 potential plastid reads (2,715,983,671 nt in total, N50 12,829 nt) and the 426,918 potential mitochondrial reads (3,928,350,102 nt, N50 12,624 nt) were separately assembled with Canu v1.7 (5). Each longest contig displayed high similarity to the chloroplast (6) or mitochondrion (7) genome sequence of *V.vinifera*. Subsequently, bandage (8) was used for confirming a correct assembly, overlapping end sequences from the circular genomes were trimmed and the start aligned to that of the grapevine reference sequences. The assemblies were polished three times with *arrow* (smrtlink-release_5.1.0.26412). The last round of polishing was carried out with the start shifted to the opposite position of the sequence.

To aid annotation, RNA was extracted from ‘Börner’ tissues using the peqGOLD Plant RNA Kit (PEQLAB) according to manufacturer’s instructions. Indexed Illumina sequencing libraries were prepared from 1000 ng total RNA according to the TruSeq RNA Sample Preparation v2 Guide. The resulting RNA-Seq libraries were pooled equimolar and sequenced 2×100 bp paired-end on a HiSeq1500.

cp_Boe (161,008 nt; GC 37.4%) and mt_Boe (755,068 nt; GC 44.3%) were annotated with the web-service GeSeq v1.66 (specific settings cp_Boe: annotate plastid IR enabled; HMMER profile search (9) enabled; reference sequence *V.vinifera* chloroplast annotation (6); MPI-MP chloroplast references enabled; specific settings mt_Boe: reference sequence *V.vinifera* mitochondrion annotation (7); settings both: tRNA annotators tRNAscan-SE v2.0 (10, 11), ARAGORN v1.2.38 (12) enabled with ‘Allow overlaps’, ‘Fix introns’) (13) that uses OGDRAW v1.3 (14, 15) to visualize the annotation (Fig. 1). RNA editing sites were determined (16) using RNA-Seq data from fife ‘Börner’ tissues. A total of 133 genes with 90 editing sites were identified for cp_Boe, encoding 85 mRNAs, 39 tRNAs, 8 rRNAs and 1 pseudogene. For mt_Boe, 172 genes with 624 editing sites were identified that encode 67 mRNAs, 38 tRNAs, 4 rRNAs and 63 pseudogenes/gene fragments. While cp_Boe confirms maternal inheritance of the chloroplast from *V*.*riparia* due to its high similarity to the chloroplast sequence from *V*.*riparia* voucher Wen 12938 (17), mt_Boe is the first mitochondrion genome sequence from *V*.*riparia* and differs from the *V.vinifera* mitochondrion (7) at 141 positions in coding regions.

**Fig. 1.**
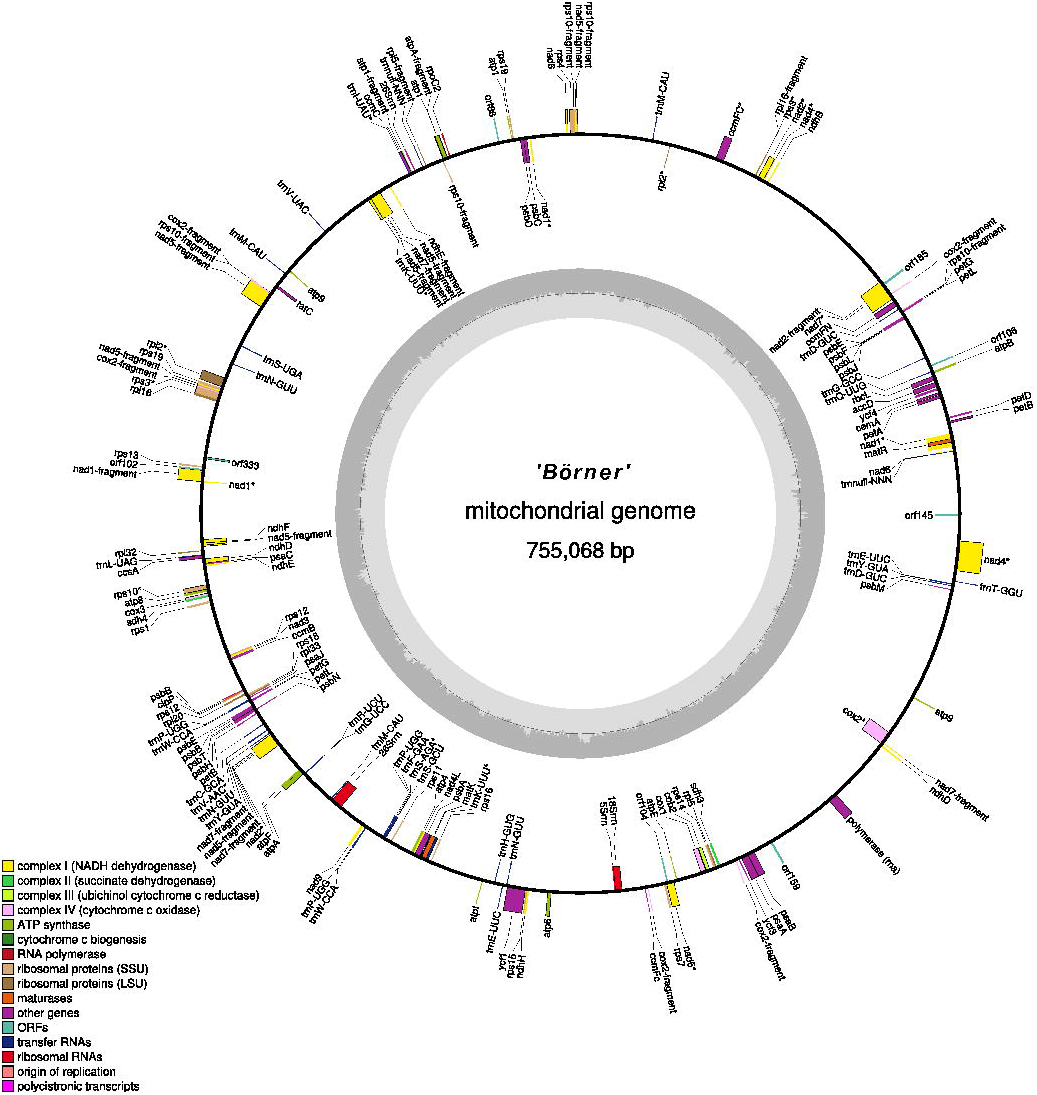
Annotation of the ‘Börner’ mitochondrial genome. Annotation was created with GeSeq and visualized with OGDRAW. Genes containing introns are marked with an asterisk (*).

## Data availability

‘Börner’ RNA-Seq reads (leaves accession no. ERR3894001, winter leaves ERR3895010, inflorescences ERR3894002, tendrils ERR3894003, roots ERR3895007), raw SMRT sequence reads (plastid ERR3610907, mitochondrion ERR3610837), the chloroplast and mitochondrion genome sequences including annotation have been deposited in GenBank/DDBJ/ENA (cp_Boe LR738917, mt_Boe, LR738918) under project number PRJEB34983. RNA editing tables, coding sequences and protein sequences of genes subject to RNA editing in edited and un-edited form are available as data publications (cpBoe_RNAedit and mtBoe_RNAedit). Data raw file and metadata were submitted according to the rules and guidelines of the COST ACTION INTEGRAPE CA17111.

## Acknowledgments

The authors thank the members of the Chair of Genetics and Genomics of Plants at Bielefeld University as well as the members of the Julius Kühn-Institute for Grapevine Breeding at Geilweilerhof for their support. The project was supported by funds of the Federal Ministry of Food and Agriculture (BMEL) based on a decision of the Parliament of the Federal Republic of Germany via the Federal Office for Agriculture and Food (BLE) under the innovation support program (project acronym MureViU, 28-1-82.066-15). We acknowledge support for the Article Processing Charge by the Deutsche Forschungsgemeinschaft and the Open Access Publication Fund of Bielefeld University.

## References

1. Ferrarini M, Moretto M, Ward JA, Surbanovski N, Stevanovic V, Giongo L, Viola R, Cavalieri D, Velasco R, Cestaro A, Sargent DJ. 2013. An evaluation of the PacBio RS platform for sequencing and de novo assembly of a chloroplast genome. BMC Genomics 14:670.

2. Stadermann KB, Weisshaar B, Holtgräwe D. 2015. SMRT sequencing only de novo assembly of the sugar beet (Beta vulgaris) chloroplast genome. BMC Bioinformatics 16:295.

3. Pucker B, Holtgräwe D, Stadermann KB, Frey K, Huettel B, Reinhardt R, Weisshaar B. 2019. A chromosome-level sequence assembly reveals the structure of the Arabidopsis thaliana Nd-1 genome and its gene set. PLoS One 14:e0216233.

4. Altschul SF, Gish W, Miller W, Myers EW, Lipman DJ. 1990. Basic local alignment search tool. Journal of Molecular Biology 215:403–410.

5. Koren S, Walenz BP, Berlin K, Miller JR, Bergman NH, Phillippy AM. 2017. Canu: scalable and accurate long-read assembly via adaptive k-mer weighting and repeat separation. Genome Research 27:722–736.

6. Jansen RK, Kaittanis C, Saski C, Lee SB, Tomkins J, Alverson AJ, Daniell H. 2006. Phylogenetic analyses of Vitis (Vitaceae) based on complete chloroplast genome sequences: effects of taxon sampling and phylogenetic methods on resolving relationships among rosids. BMC Evolutionary Biology 6:32.

7. Goremykin VV, Salamini F, Velasco R, Viola R. 2009. Mitochondrial DNA of Vitis vinifera and the issue of rampant horizontal gene transfer. Molecular Biology and Evolution 26:99–110.

8. Wick RR, Schultz MB, Zobel J, Holt KE. 2015. Bandage: interactive visualization of de novo genome assemblies. Bioinformatics 31:3350–2.

9. Wheeler TJ, Eddy SR. 2013. nhmmer: DNA homology search with profile HMMs. Bioinformatics 29:2487–2489.

10. Chan PP, Lowe TM. 2019. tRNAscan-SE: Searching for tRNA genes in genomic sequences, p 1-14. In Kollmar M (ed), Gene Prediction: Methods and Protocols, 2019/04/26 ed, vol 1962. Springer New York, New York.

11. Lowe TM, Eddy SR. 1997. tRNAscan-SE: a program for improved detection of transfer RNA genes in genomic sequence. Nucleic Acids Research 25:955–964.

12. Laslett D, Canback B. 2004. ARAGORN, a program to detect tRNA genes and tmRNA genes in nucleotide sequences. Nucleic Acids Research 32:11–16.

13. Tillich M, Lehwark P, Pellizzer T, Ulbricht-Jones ES, Fischer A, Bock R, Greiner S. 2017. GeSeq - versatile and accurate annotation of organelle genomes. Nucleic Acids Research 45:W6–W11.

14. Lohse M, Drechsel O, Kahlau S, Bock R. 2013. OrganellarGenomeDRAW--a suite of tools for generating physical maps of plastid and mitochondrial genomes and visualizing expression data sets. Nucleic Acids Research 41:W575–581.

15. Lohse M, Drechsel O, Bock R. 2007. OrganellarGenomeDRAW (OGDRAW): a tool for the easy generation of high-quality custom graphical maps of plastid and mitochondrial genomes. Current Genetics 52:267–274.

16. Brenner WG, Mader M, Muller NA, Hoenicka H, Schroeder H, Zorn I, Fladung M, Kersten B. 2019. High Level of Conservation of Mitochondrial RNA Editing Sites Among Four Populus Species. G3 (Bethesda) 9:709–717.

17. Wen J, Harris AJ, Kalburg Y, Zhang N, Xu Y, Zheng W, Ickert-Bond SM, Johnson G, Zimmer EA. 2018. Chloroplast phylogenomics of the New World grape species (Vitis, Vitaceae). Journal of Systematics and Evolution 56:297–308.

